# A hybrid unsupervised approach for accurate short read clustering and barcoded sample demultiplexing in nanopore sequencing

**DOI:** 10.1101/2022.04.13.488186

**Authors:** Renmin Han, Junhai Qi, Yang Xue, Xiujuan Sun, Fa Zhang, Xin Gao, Guojun Li

## Abstract

Short nucleic acid sequences are usually attached as DNA barcodes for multiple sample sequencing and single cell protocols, which enables Oxford Nanopore sequencing to sequence multiple barcoded DNA samples on a single flow cell. However, due to the high base-calling error, short reads in Nanopore sequencing are difficult to be accurately identified by traditional tools. Here, we propose a hybrid unsupervised approach for the accurate clustering of short reads and demultiplexing of barcoded samples in Nanopore sequencing. In our approach, both the nucleic base information translated from base-calling and the raw current signal directly outputted by the flow cell are utilized. A GPU-supported parallelization strategy is proposed to ensure the runtime of our hybrid clustering. Comprehensive experiments demonstrate that our approach outperforms all the traditional unsupervised tools in short read clustering, and achieves comparable accuracy in barcoded sample demultiplexing compared with the learning-based methods.

## 1 Background

A barcode is a very short nucleotide sequence attached at the 3’- or 5’-end of a DNA sequence to state where the sequence comes. By incorporating a unique barcode into the library of DNA molecules, multiple DNA libraries are able to be sequenced simultaneously [1]. Usually, the length of these short nucleotide sequences ranges from 20 to 140 bp. Clustering or classifying the reads into bins based on these short nucleic acid fragements is the first step in high-throughput sequencing techniques like multiple sample sequencing and single cell protocols [2, 3]. Specifically, the barcoding technique has recently been introduced to Oxford Nanopore devices to sequence multiple barcoded DNA samples on a single flow cell [4, 5].

Oxford Nanopore sequencing is a rapidly developing technology that enables ultra-long sequencing in real time at low-cost. The key innovation of nanopore sequencing is the direct measurement of the electrical current signal (denoted as the *raw signal*) when a single-strand DNA passes through the nanopore. These raw signals are transferred to nucleic acid bases by base-calling for further analysis [6, 7, 8]. The translation from raw current signals to reads may introduce significant base-calling errors. Specifically, considering a 40-bp barcode and a base-calling system with 10% error, the possibility that a sequenced barcode is completely correct is 0.9^40^ ≈ 0.014, which can badly hamper the downstream analyses [9]. Especially, because of the high base-calling error, the Unique Molecular Identifiers (UMI) technique, which is shorter nucleotide sequence added to sequencing libraries to identify PCR duplicates, is rarely used in Nanopore sequencing [10, 11].

A number of methods have been devised to group biological sequences that are related. In early 2001, a tool named CD-HIT [12] is proposed for the clustering of a large number of sequences, based on pairwise alignment and greedy strategy. Later, improved methods [13, 14] of CD-HIT are also devised to cope with the next-generation sequencing data. Inspired by CD-HIT, DNACLUST [15] is proposed for taxonomic profiling. Recently, the mean shift algorithm has also been introduced by MeShClust [16] to reduce the side effect of parameter dependency in the greedy strategy. On the other hand, alignment-free similarity measures [17, 18, 19, 20] have been utilized in sequence clustering [21, 22], by mapping DNA sequences into feature vectors. Furthermore, clustering tools have also been devised for specified purpose, e.g. the Starcode [23] and Bartender [24]. However, all of these methods could only utilize the information of base-called reads in Nanopore sequencing.

On the contrary, the raw current signal contains much more information compared with the base-called reads. In practice, the frequency of the electrical current measurements is 7-9 times higher than the passing speed of the DNA sequence, which makes the raw current signal to contain ∼8× redundant information than the base-called read. Except for signal-level polishing [25], efforts have been made to utilize raw signal for targeted sequencing [26, 8, 27], variant identification [28, 29] and methylation detection [30, 31, 32]. Recently, the raw current signal has also been utilized in ONT barcode demultiplexing and achieved good results, by training a deep neural network as barcode’s raw signal classifier [33, 34]. Here, the demultiplexing is carried out as a supervised machine learning task with the classifier trained under a large human-labelled dataset. However, a problem with the supervised-learning-based classification is that the performance of these methods heavily depends on the training dataset.

In this paper, we first demonstrate that, for Nanopore sequencing, signal-similarity (dynamic time warping distance) based clustering performs much better than the base-space clustering in various criteria (Supplementary Material S1), though the computation of pair-wise signal similarity is computationally expensive (Supplementary Material S2). Consequently, we propose a hybrid unsupervised approach for the accurate clustering of short reads and demultiplexing of barcoded samples in Nanopore sequencing. Our approach utilizes both the base-called nucleic base information and the raw current signal, in which the base-called nucleic bases are used to generate initial clustering and representation sequences, while the raw signals are used for cluster merging and refinement (Figure 1 gives an example). A checking mechanism is built to make sure that the good sequences are reserved and correctly grouped. Comprehensive experiments demonstrate that our approach is highly effective in short read clustering, outperforming all the traditional unsupervised tools, with an accuracy of ∼ 95%. Moreover, parallel algorithms are designed to take advantage of the widely-used GPU cards. With a standard GPU card, our approach could achieve the clustering analysis of 1 GB data within 5 minutes.

**Figure 1.**
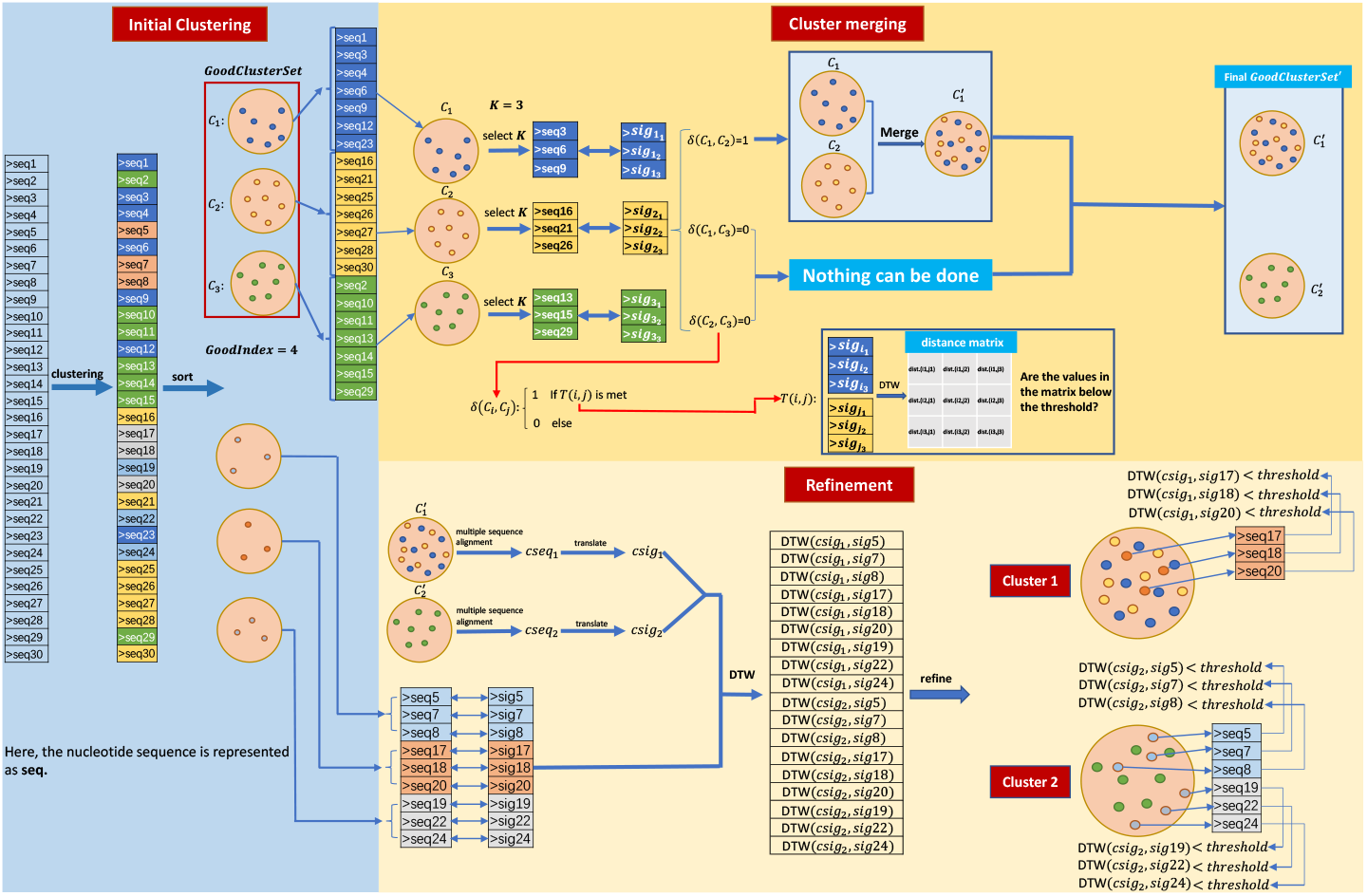
An example of clustering 30 sequences using our hybrid approach. Because there are no ultra-short sequences among the 30 sequences, no sequence filtered in the initial clustering.

## 2 Results

we designed an unsupervised hybrid approach to achieve accurate and efficient short read clustering for Nanopore sequencing, in which the nucleobase-based greedy algorithm is utilized to obtain initial clusters (Initial clustering), and the raw signal information is measured to guide the continuously optimization and refinement of clustering results (Cluster merging and Cluster refinement). GPU acceleration based on CUDA technique is utilized in our hybrid clustering (GPU-accelerated DTW). The detailed implementation is explicated in Methods Section.

### 2.1 Datasets

#### Simulated datasets

A set of synthetic datasets with different configurations are generated. Here, we first generate a set of random barcodes, and then produce a number of these barcodes’ copies as well as their raw signals by DeepSimulator. The configuration of synthetic dataset includes the following three points:

- The length of barcode (nucleotide sequence length).
- The number of clusters within a dataset.
- The number of sequences for the whole dataset.

Finally, we construct 12 simulated datasets. The details of these datasets are provided in Supplementary Material S4.

#### Real-world datasets

The real-world dataset came from eight R9.4 flow cells and six R9.5 flow cells, all sequenced with the EXP-NBD103 barcoding kit (denoted as amplicon library) [33]. In amplicon library, we use Edlib [35] to locate the fixed region of the barcode and segment the barcode read, and use Semi-Global Dynamic Time Warping [36] to extract the corresponding raw signal of these barcodes. Finally, we obtain 120,947 barcode reads (signals).

### 2.2 Run scripts

DNACLSUT fails to cope with dataset large than 10,000 sequences. Therefore, here we mainly compare our hybrid clustering with CD-HIT, UCLUST, MeShClust. The command line options for these three clustering tools are listed as follows:

1. CD-HIT: ./cd-hit-est -i infile.fasta -o outfile.fasta -c indentity
2. MeShClust: ./meshclust infile.fasta –id identity -output outfile.fasta
3. UCLSUT: ./usearch -cluster fast infile.fasta -id identity -clusters output

All the experiments were run on an Ubuntu 18.04 system with Intel(R) Core(TM) i9-10980XE (18 cores), 128 Gb memory, and an NVIDIA RTX3080 card.

### 2.3 Evaluation on synthetic datasets

Six synthetic datasets with different barcode lengths, numbers of clusters and data sizes are selected to demonstrate the performance of hybrid clustering. Table 1 describes the details of the six selected datasets.

**Table 1.** Summaries of the details about the selected synthetic datasets

| Dataset | Barcode length | Number of clusters | Data size |
| --- | --- | --- | --- |
| $S_1$ | 45bp | 20 | 50000 |
| $S_2$ | 70bp | 20 | 50000 |
| $S_3$ | 95bp | 20 | 50000 |
| $S_4$ | 45bp | 50 | 50000 |
| $S_5$ | 70bp | 50 | 50000 |
| $S_6$ | 95bp | 50 | 50000 |

Table 2-7 summarize the experimental results of different clustering methods on these synthetic datasets, where the indexes AMI, FMI, ACC, HOMO, COMP and runtime are adapted for the performance evaluation.

**Table 2.**
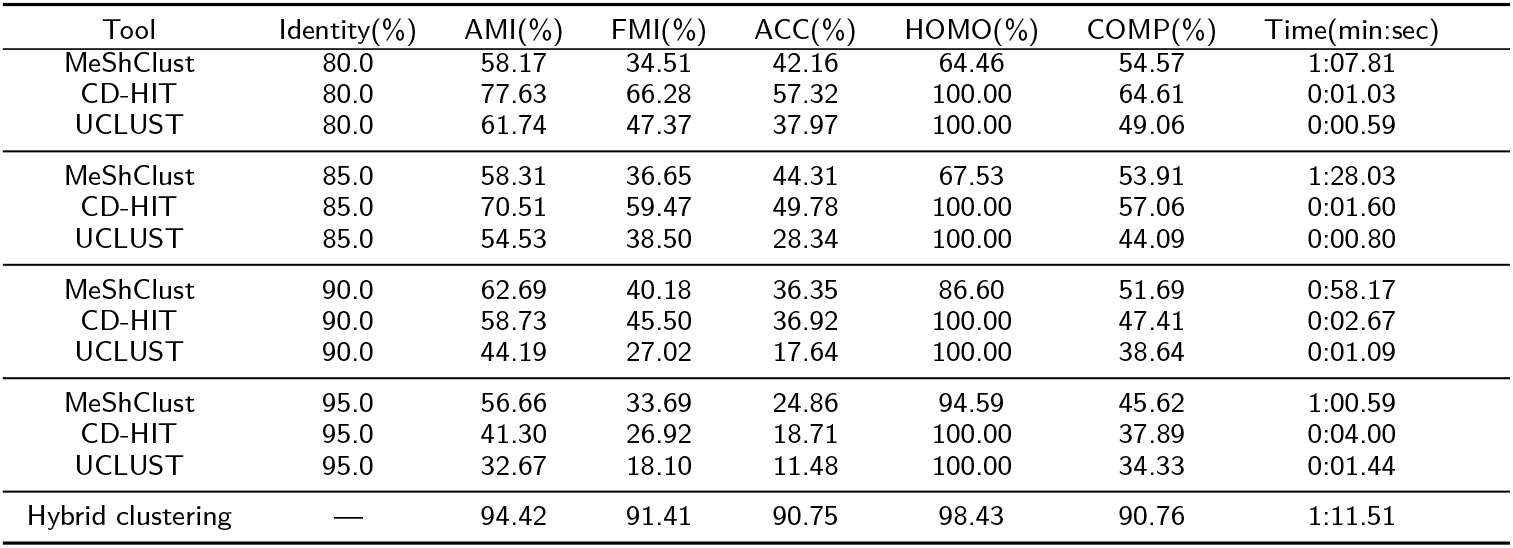
Performance comparison of the different clustering methods on the dataset *S*_1_

Judging from the experimental results, it can be found that CD-HIT and UCLUST are able to guarantee high homogeneity (HOMO index = 100%) under various situations. The high homogeneity is reasonable because CD-HIT and UCLUST are designed to maintain the consistency of the elements within a cluster. UCLUST is the fastest and CD-HIT is the second fastest in clustering speed, because of the utility of non-alignment technique. When the sequence length is very short, MeShClust behaves the poorest within the four clustering methods, as shown in Table 2.

Drown from the six synthetic datasets, we can make the following key conclusions:

- CD-HIT and UCLUST are the fastest and able to guarantee high homogeneity (HOMO more than 98%) of the results.
- For barcodes with short sequence length (*S*_1_ and *S*_4_), the clustering performance of MeShClust is very poor. With the increase of barcode length, the clustering performance of MeShClust is significantly improved.
- The clustering results of *S*_1_ to *S*_3_ (Table 2∼4) and *S*_4_ to *S*_6_ (Table 5∼7) demonstrate that the performance of clustering tools depends on the length of barcode sequence, and longer sequence results in better clustering result.
- Although the clustering speed of the hybrid clustering is relatively slow, its performance is significantly better than the other methods. Except for the homogeneity (HOMO) metric, the hybrid clustering gains almost 20% performance improvement in all the other indexes.
- The length of the sequence will affect the speed of hybrid clustering, but this does not mean that the longer the sequence, the slower the clustering speed as shown in Table 6 and Table 7. The reason for this phenomenon may be that the longer the sequence, the better the initial clustering performance, which reduces the time for cluster merging and refinement in hybrid clustering.
- The speed of hybrid clustering is affected by the number of clusters residing in the dataset, as the result comparison of *S*_1_ and *S*_4_, *S*_2_ and *S*_5_, and *S*_3_ and *S*_6_. This is caused by the fact that the number of clusters determines the number of DTW distance that should be calculated in the cluster merging phase.

**Table 3.** Performance comparison of the different clustering methods on the dataset *S*_2_

| Tool | Identity(%) | AMI(%) | FMI(%) | ACC(%) | HOMO(%) | COMP(%) | Time(min:sec) |
| --- | --- | --- | --- | --- | --- | --- | --- |
| MeShClust | 80.0 | 67.10 | 50.96 | 57.66 | 70.45 | 64.98 | 0:55.28 |
| CD-HIT | 80.0 | 81.89 | 71.30 | 65.57 | 100.00 | 70.02 | 0:01.40 |
| UCLUST | 80.0 | 63.74 | 45.82 | 35.77 | 100.00 | 50.27 | 0:00.96 |
| MeShClust | 85.0 | 67.63 | 52.12 | 52.67 | 83.91 | 58.61 | 1:12.78 |
| CD-HIT | 85.0 | 73.60 | 60.10 | 52.22 | 100.00 | 60.09 | 0:02.36 |
| UCLUST | 85.0 | 53.73 | 32.44 | 22.27 | 100.00 | 43.06 | 0:01.35 |
| MeShClust | 90.0 | 61.96 | 50.08 | 41.15 | 92.98 | 50.83 | 1:00.49 |
| CD-HIT | 90.0 | 60.24 | 42.76 | 33.59 | 100.00 | 48.08 | 0:04.63 |
| UCLUST | 90.0 | 40.88 | 20.52 | 13.63 | 100.00 | 36.89 | 0:01.91 |
| MeShClust | 95.0 | 59.67 | 49.57 | 44.31 | 96.26 | 49.03 | 1:56.98 |
| CD-HIT | 95.0 | 40.52 | 24.03 | 16.17 | 100.00 | 37.37 | 0:07.25 |
| UCLUST | 95.0 | 26.29 | 12.46 | 7.34 | 100.00 | 32.38 | 0:02.44 |
| Hybrid clustering | — | 96.82 | 95.61 | 95.46 | 100.00 | 93.86 | 2:00.98 |

**Table 4.** Performance comparison of the different clustering methods on the dataset *S*_3_

| Tool | Identity(%) | AMI(%) | FMI(%) | ACC(%) | HOMO(%) | COMP(%) | Time(min:sec) |
| --- | --- | --- | --- | --- | --- | --- | --- |
| MeShClust | 80.0 | 85.06 | 83.40 | 86.24 | 93.44 | 79.06 | 1:33.12 |
| CD-HIT | 80.0 | 88.70 | 83.32 | 78.95 | 100.00 | 79.99 | 0:01.65 |
| UCLUST | 80.0 | 68.75 | 54.50 | 44.48 | 100.00 | 55.08 | 0:01.29 |
| MeShClust | 85.0 | 79.99 | 81.27 | 79.19 | 96.50 | 70.48 | 1:58.35 |
| CD-HIT | 85.0 | 81.22 | 74.83 | 67.45 | 100.00 | 69.48 | 0:03.20 |
| UCLUST | 85.0 | 58.44 | 44.18 | 32.51 | 100.00 | 46.91 | 0:01.79 |
| MeShClust | 90.0 | 66.37 | 65.43 | 61.96 | 94.12 | 56.85 | 2:01.84 |
| CD-HIT | 90.0 | 67.86 | 60.14 | 52.22 | 100.00 | 55.24 | 0:06.92 |
| UCLUST | 90.0 | 44.36 | 26.50 | 18.00 | 100.00 | 38.76 | 0:02.46 |
| MeShClust | 95.0 | 57.65 | 58.07 | 45.25 | 99.47 | 48.83 | 1:07.26 |
| CD-HIT | 95.0 | 44.53 | 30.26 | 21.45 | 100.00 | 39.50 | 0:04.00 |
| UCLUST | 95.0 | 26.93 | 14.09 | 8.60 | 100.00 | 32.67 | 0:01.44 |
| Hybrid clustering | — | 97.74 | 96.72 | 96.42 | 100.00 | 95.59 | 1:56.77 |

**Table 5.**
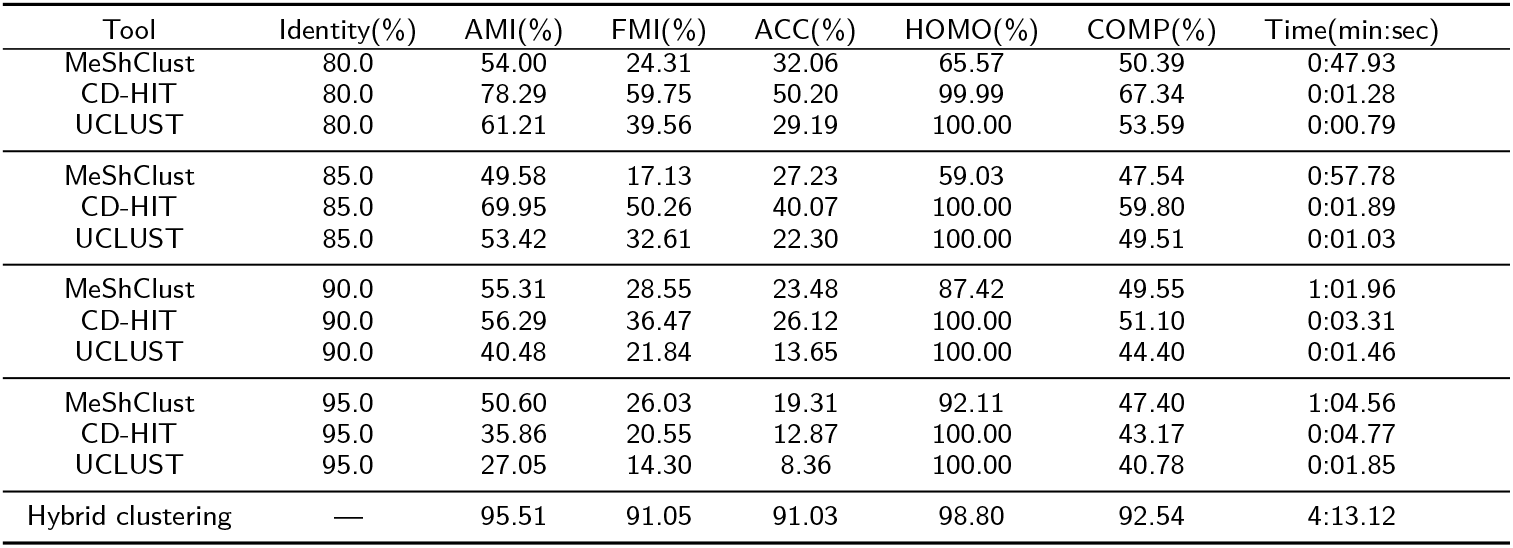
Performance comparison of the different clustering methods on the dataset *S*_4_

| Tool | Identity(%) | AMI(%) | FMI(%) | ACC(%) | HOMO(%) | COMP(%) | Time(min:sec) |
| --- | --- | --- | --- | --- | --- | --- | --- |
| MeShClust | 80.0 | 54.00 | 24.31 | 32.06 | 65.57 | 50.39 | 0:47.93 |
| CD-HIT | 80.0 | 78.29 | 59.75 | 50.20 | 99.99 | 67.34 | 0:01.28 |
| UCLUST | 80.0 | 61.21 | 39.56 | 29.19 | 100.00 | 53.59 | 0:00.79 |
| MeShClust | 85.0 | 49.58 | 17.13 | 27.23 | 59.03 | 47.54 | 0:57.78 |
| CD-HIT | 85.0 | 69.95 | 50.26 | 40.07 | 100.00 | 59.80 | 0:01.89 |
| UCLUST | 85.0 | 53.42 | 32.61 | 22.30 | 100.00 | 49.51 | 0:01.03 |
| MeShClust | 90.0 | 55.31 | 28.55 | 23.48 | 87.42 | 49.55 | 1:01.96 |
| CD-HIT | 90.0 | 56.29 | 36.47 | 26.12 | 100.00 | 51.10 | 0:03.31 |
| UCLUST | 90.0 | 40.48 | 21.84 | 13.65 | 100.00 | 44.40 | 0:01.46 |
| MeShClust | 95.0 | 50.60 | 26.03 | 19.31 | 92.11 | 47.40 | 1:04.56 |
| CD-HIT | 95.0 | 35.86 | 20.55 | 12.87 | 100.00 | 43.17 | 0:04.77 |
| UCLUST | 95.0 | 27.05 | 14.30 | 8.36 | 100.00 | 40.78 | 0:01.85 |
| Hybrid clustering | — | 95.51 | 91.05 | 91.03 | 98.80 | 92.54 | 4:13.12 |

**Table 6.** Performance comparison of the different clustering methods on the dataset *S*_5_

| Tool | Identity(%) | AMI(%) | FMI(%) | ACC(%) | HOMO(%) | COMP(%) | Time(min:sec) |
| --- | --- | --- | --- | --- | --- | --- | --- |
| MeShClust | 80.0 | 68.39 | 47.68 | 52.91 | 82.86 | 62.65 | 1:03.37 |
| CD-HIT | 80.0 | 85.39 | 71.10 | 64.08 | 100.00 | 75.94 | 0:01.66 |
| UCLUST | 80.0 | 70.56 | 55.00 | 43.23 | 100.00 | 60.60 | 0:01.13 |
| MeShClust | 85.0 | 62.40 | 36.39 | 46.49 | 82.04 | 57.61 | 1:43.73 |
| CD-HIT | 85.0 | 77.87 | 60.19 | 51.56 | 100.00 | 67.23 | 0:02.76 |
| UCLUST | 85.0 | 60.71 | 44.25 | 32.51 | 100.00 | 53.79 | 0:01.56 |
| MeShClust | 90.0 | 54.90 | 24.28 | 33.06 | 73.30 | 52.13 | 1:55.66 |
| CD-HIT | 90.0 | 65.62 | 48.89 | 39.54 | 100.00 | 57.13 | 0:04.93 |
| UCLUST | 90.0 | 46.99 | 30.48 | 20.19 | 100.00 | 47.03 | 0:02.12 |
| MeShClust | 95.0 | 53.89 | 24.53 | 31.02 | 82.26 | 51.23 | 1:06.23 |
| CD-HIT | 95.0 | 45.10 | 28.76 | 20.16 | 100.00 | 46.40 | 0:07.29 |
| UCLUST | 95.0 | 30.91 | 18.12 | 10.80 | 100.00 | 41.81 | 0:02.50 |
| Hybrid clustering | — | 97.05 | 93.70 | 92.40 | 99.99 | 94.36 | 5:41.25 |

**Table 7.** Performance comparison of the different clustering methods on the dataset *S*_6_

| Tool | Identity(%) | AMI(%) | FMI(%) | ACC(%) | HOMO(%) | COMP(%) | Time(min:sec) |
| --- | --- | --- | --- | --- | --- | --- | --- |
| MeShClust | 80.0 | 87.66 | 79.22 | 83.36 | 93.81 | 83.40 | 0:49.57 |
| CD-HIT | 80.0 | 92.46 | 86.29 | 82.23 | 100.00 | 86.44 | 0:01.58 |
| UCLUST | 80.0 | 78.99 | 67.27 | 57.71 | 100.00 | 68.73 | 0:01.27 |
| MeShClust | 85.0 | 83.44 | 76.77 | 77.05 | 95.24 | 76.69 | 1:01.72 |
| CD-HIT | 85.0 | 86.30 | 77.45 | 72.41 | 100.00 | 77.51 | 0:02.80 |
| UCLUST | 85.0 | 69.14 | 54.09 | 44.34 | 100.00 | 59.75 | 0:01.73 |
| MeShClust | 90.0 | 76.70 | 71.47 | 68.35 | 96.18 | 68.72 | 1:03.75 |
| CD-HIT | 90.0 | 74.23 | 61.01 | 52.30 | 100.00 | 64.25 | 0:05.54 |
| UCLUST | 90.0 | 53.36 | 34.91 | 25.64 | 100.00 | 49.81 | 0:02.39 |
| MeShClust | 95.0 | 70.43 | 64.56 | 56.08 | 98.07 | 62.35 | 1:05.74 |
| CD-HIT | 95.0 | 52.12 | 32.98 | 24.42 | 100.00 | 49.34 | 0:09.23 |
| UCLUST | 95.0 | 31.74 | 15.71 | 9.68 | 100.00 | 41.86 | 0:03.07 |
| Hybrid clustering | — | 97.80 | 95.72 | 95.26 | 99.99 | 95.75 | 4:52.07 |

In general, due to the base-calling error, the traditional clustering tools such as CD-HIT, UCLUST and MeShClust could not get good clustering results in the analysis of short nanopore reads. The initial clustering of our method guarantees the extremely high homogeneity within the clusters, and the cluster merging and refinement guarantee the high accuracy and completeness of the clustering, which greatly reduces the influence of base-calling error. The results show that our method produces very good clustering results, and the results produced by traditional clustering tools cannot compete with us. More benchmarking results are provided in Supplementary Material S4 (Table S10-Table S15).

### 2.4 Performance analysis of different stages

As introduced in previous sections, the hybrid clustering algorithm is composed by three stages, i.e., initial clustering, cluster merge and cluster refinement. In this section, we analyze the detailed contributions of different stages in hybrid clustering and their time cost.

First, we analyze the change of cluster accuracy of these different stages. As shown in Figure 2, after the initial clustering, the clustering result is not so good. With the completion of the merging phase, the clustering performance has been greatly improved. After the refinement phase, the clustering result is further improved. The change of index values in Figure 2 clearly show the effectiveness of the three-stage solution in our hybrid clustering algorithm. Especially, the signal based cluster merging and refinement contributes a lot to the accuracy improvement in clustering result.

**Figure 2.**
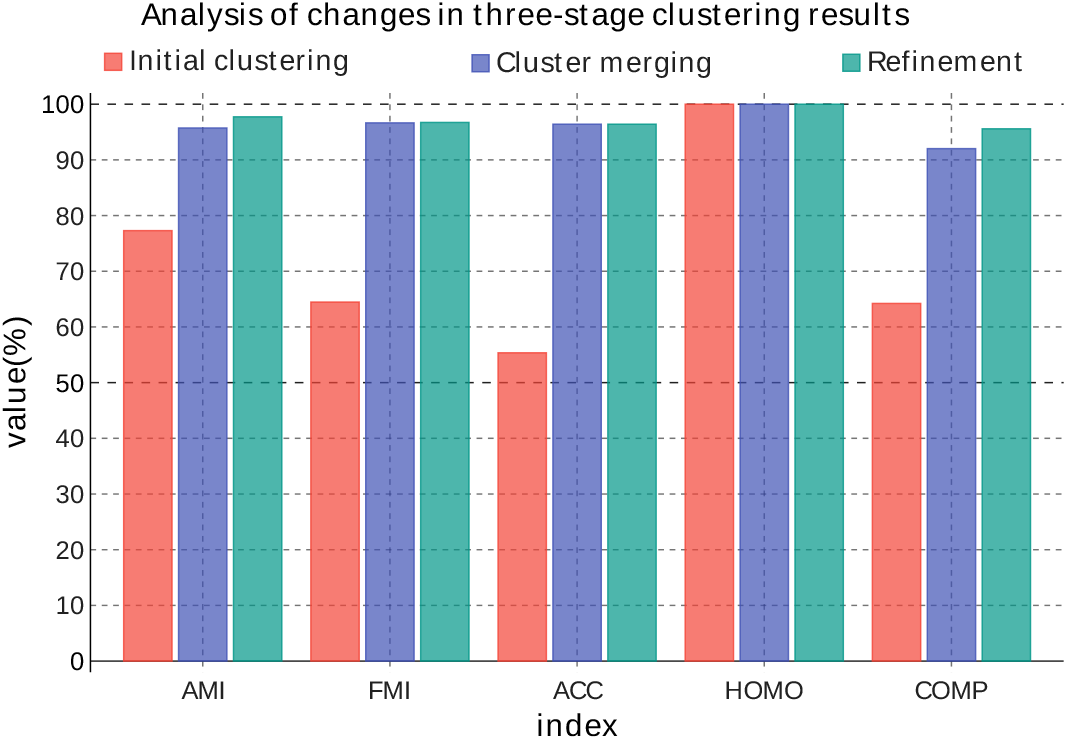
The clustering result of hybrid clustering at different stages on dataset *S*_3_. The results after initial clustering, cluster merging and cluster refinement is labeled by red, purple and green, respectively.

Then, we analyze the time cost of these different stages. Figure 3 shows the runtime of the three stages for the clustering of *S*_1_ in pie chart. As shown in Figure 3A, the hybrid clustering algorithm spends the most time in the cluster merging stage. It is reasonable because a huge number of DTW distance comparison and multiple sequence alignment algorithm are calculated in the cluster merging, both of which are relatively time-consuming. The detailed time cost percentage of DTW distance comparison between clusters and the generation of consensus sequence are shown in Figure 3B. Here, it can be found that the generation of consensus sequence costs more than the DTW distance comparison. This could be true because we have accelerated the calculation of DTW by GPU but the generation of consensus sequence is still supported by CPU.

**Figure 3.**
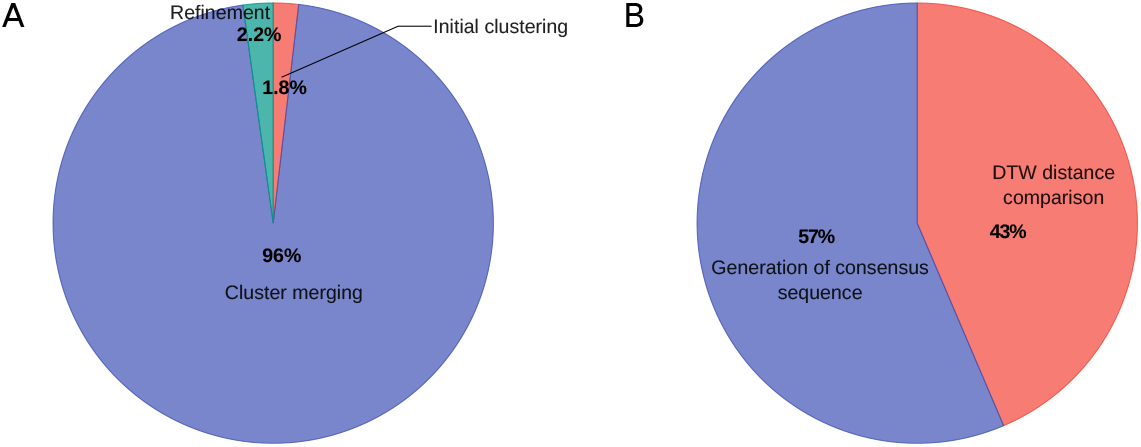
Pie chart of different stages’ runtime percentage of the algorithm on *S*_1_. (A) the three stages’ runtime percentage. (B) The runtime percentage of the DTW distance comparison and the generation of consensus sequence in cluster merging.

### 2.5 Speedup of GPU-accelerated DTW

As described in previous section, numerous DTW distance are calculated in our algorithm but we still achieve relatively small time cost. Here, we would like to show the benefits of GPU acceleration in DTW distance calculation.

In order to show the overall acceleration effect, we generate a large amount of time series as test data, whose details are shown in Tabel 8. We compare the CUDA implementation of DTW with CPU single-threaded method and CPU multi-threaded method. The DTW method realized by CUDA is equivalent to the original DTW in the mathematical model, which can guarantee its correctness. The CPU single-threaded method is a naive DTW algorithm. In the CPU multi-threaded approach, each CPU thread is responsible for calculating the DTW distance of a pair of time series, while the single CPU thread still uses the traditional method for calculation. Figure 4 clearly shows the time spent by different approaches in logarithmic scale. As shown in Figure 4, the CUDA accelerated DTW is at least three orders of magnitude faster than the traditional single-threaded DTW, and two orders of magnitude faster than the 30-threaded DTW.

**Figure 4.**
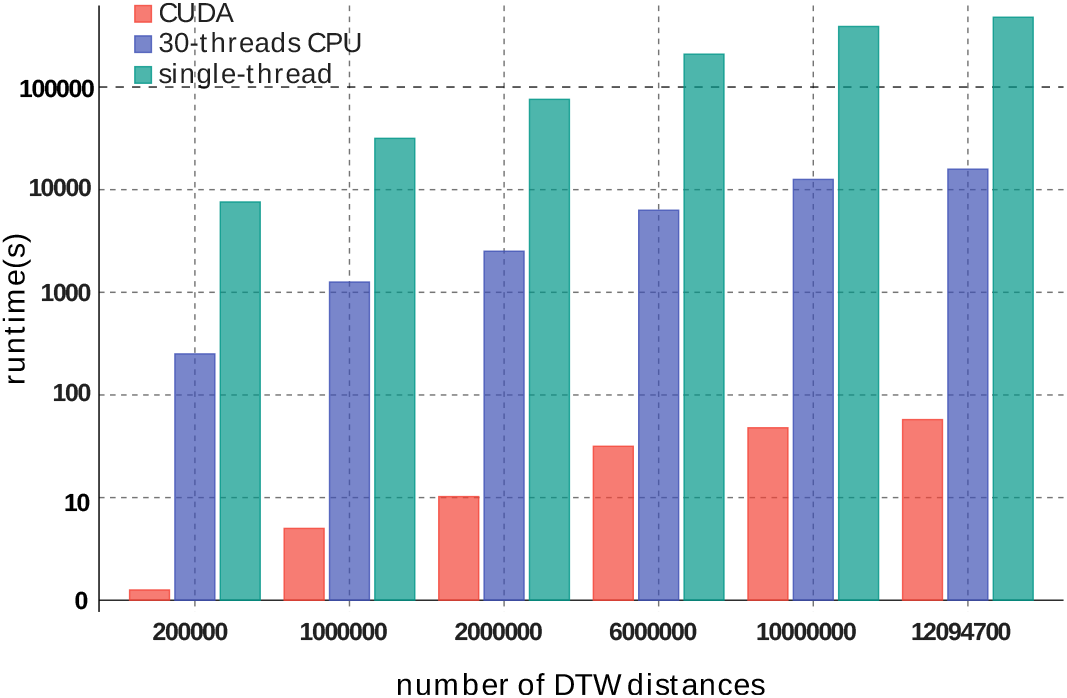
The time comparison between different acceleration strategy with different data size.

In addition, we evaluate the DTW acceleration ratio of different lengths by simulated nanopore signals generated from DeepSimulator, where The length of the current signal is approximately 8 times of that of the corresponding DNA template. Figure 5 shows the change of acceleration ratio with different sequence lengths, where the acceleration ratio for single DTW calculation ranges from 14× to 22×, increasing with the lengthening of sequences. Furthermore, as introduced in previous section, a block-wise acceleration strategy is proposed to fully utilize the advantage of GPU blocks, which enables the launch of million threads of DTW calculation simultaneously. In Supplementary Table S1, we have shown that it takes about 1100 minutes to calculate the DTW distance matrix of 2000 × 2000. As a comparison, by applying the CUDA acceleration strategy, the time cost of the DTW distance matrix calculation can be reduced to 4 seconds.

**Figure 5.**
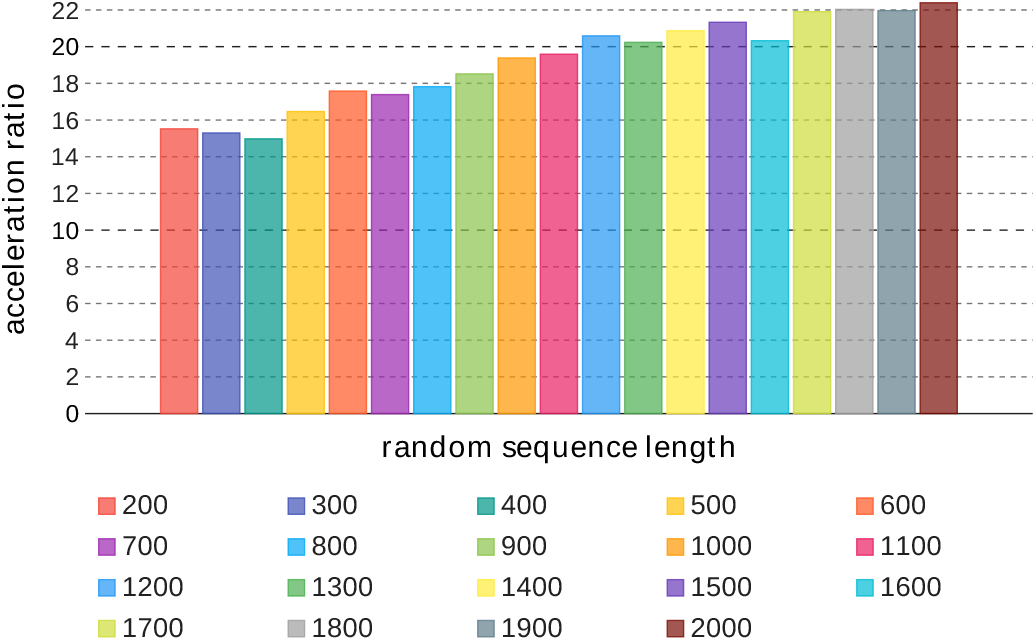
Acceleration ratio for random sequences with different lengths., where the acceleration ratio = (runtime of single-thread DTW)/(runtime of single-thread CUDA).

### 2.6 Runtime analysis of the whole algorithm

As discussed in previous section, our algorithm consists of three stages and the calculation of DTW distance is GPU-accelerated. Here, we would like to further analyze the overall time complexity of the hybrid clustering algorithm.

We simulated a number of datasets by DeepSimulator to test the time cost of our algorithm under different sequence lengths, dataset sizes and numbers of clusters, whose results are summarized in Figure 6. Figure 6A shows that the runtime of our algorithm is not simply correlated with the sequence length. When the sequence length is 55-bp, the total time cost of our algorithm is smaller than the one with 45-bp sequence. In addition, the time costs for datasets with 75-bp, 85-bp and 95-bp sequence length are almost the same. In fact, the accuracy of initial clustering could benefit from longer sequence, which shortens the runtime of further cluster merging and refinement. Figure 6B and 6C shows that the runtime of our algorithm linearly increases with the increment of dataset size and number of clusters, which ensures an acceptable time cost even when the size of dataset is relatively large. In practice, our method can complete barcode clustering efficiently.

**Figure 6.**
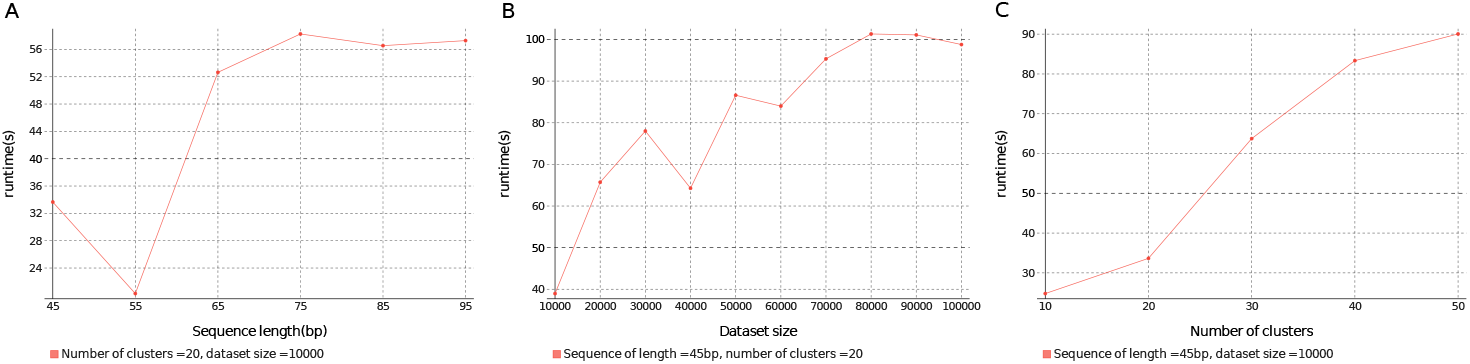
The runtime of hybrid clustering on (A) a dataset with 10,000 sequences and 20 clusters when the length of the sequences changes, (B) a dataset with about 45bp long sequences and 20 clusters when the size changes, and (C) a dataset with long sequences when the number of clusters changes.

### 2.7 Evaluation on real-world dataset

The real-world dataset is downloaded from [33], composed of 12 classes of nanopore barcodes, with ∼40 base pairs for each barcode. For the real-world barcode sequences, the first 8 positions of nucleobase and the last 8 positions of nucleobase are same to each other. Thus, the base-calling error may translate two identical nanopore barcode into different nucleobase reads, which greatly hampers the correctness of clustering and classification.

Table 9 summarizes the experimental results of different clustering methods on the real-world dataset. As shown in Table 9, the performance of MeShClust on the real-world dataset is very poor at all identities, failing to guarantee even the homogeneity of clustering. At each identity, the performance of the CD-HIT is slightly better than that of UCLUST, while UCLUST can always guarantee higher HOMO. The performance of hybrid clustering algorithm on the real-world dataset is much better than that of the other classic clustering tools, with 95.36% accuracy and 93.18% completeness. In terms of clustering efficiency, UCLUST and CDHIT still maintain a clear advantage, while the hybrid clustering algorithm can also complete the clustering of about one hundred thousand sequences within very short time. By fully utilizing the raw signal information, the hybrid clustering algorithm can cope with the challenge of base-calling error well, outperforming the classic ‘base-space’ clustering tools. Especially, our algorithm has finished the clustering within 1 minute, which is also very efficient.

**Table 8.** Three different kinds of time series used for the comparison of different DTW’s implementation, where all the time series are with a length of 1300. “Random” means time series of random walk. “Simulator” means current signals generated by DeepSimulator. “Amplicon library” means the real data downloaded from [33]. Here, we divide the data into two groups, and the time series within each group will compare with each other. For example, in the first random dataset, 200 *×* 1000 means that the first group has 200 time series, the second group has 1000 time series, and the number of DTW comparison is 200 *×* 1000 = 200000.

| Source | size | DTW numbers |
| --- | --- | --- |
| Random | 200×1000 | 200000 |
| Random | 1000×1000 | 1000000 |
| Simulator | 200×10000 | 2000000 |
| Random | 200×30000 | 6000000 |
| Random | 1000×10000 | 10000000 |
| Amplicon library | 100×120947 | 12094700 |

**Table 9.** Performance comparison of the different clustering methods on the real-world dataset.

| Tool | Identity(%) | AMI(%) | FMI(%) | ACC(%) | HOMO(%) | COMP(%) | Time(min:sec) |
| --- | --- | --- | --- | --- | --- | --- | --- |
| MeShClust | 80.0 | 0.20 | 30.37 | 15.93 | 0.32 | 19.02 | 0:51.35 |
| CD-HIT | 80.0 | 77.86 | 71.68 | 64.68 | 96.31 | 65.57 | 0:02.36 |
| UCLUST | 80.0 | 56.86 | 35.70 | 20.96 | 99.96 | 41.24 | 0:02.2 |
| MeShClust | 85.0 | 4.63 | 29.53 | 17.13 | 3.14 | 25.76 | 1:31.95 |
| CD-HIT | 85.0 | 70.21 | 56.74 | 50.29 | 99.77 | 54.71 | 0:03.14 |
| UCLUST | 85.0 | 50.08 | 27.07 | 16.35 | 99.97 | 36.12 | 0:01.73 |
| MeShClust | 90.0 | 27.06 | 23.52 | 23.02 | 24.44 | 34.50 | 3:02.26 |
| CD-HIT | 90.0 | 56.88 | 34.81 | 27.64 | 99.96 | 41.60 | 0:07.89 |
| UCLUST | 90.0 | 40.92 | 18.21 | 12.35 | 99.98 | 30.93 | 0:02.79 |
| MeShClust | 95.0 | 26.54 | 21.75 | 18.98 | 29.42 | 30.36 | 7:31.98 |
| CD-HIT | 95.0 | 39.71 | 18.30 | 13.56 | 99.98 | 30.72 | 0:21.57 |
| UCLUST | 95.0 | 31.10 | 13.39 | 9.63 | 99.99 | 27.32 | 0:04.32 |
| Hybrid clustering | — | 94.74 | 94.29 | 95.36 | 95.96 | 93.18 | 0:45.86 |

## 3 Discussion

Nanopore sequencing produces two types of data, i.e., the raw current signals and base-called reads. For barcode sequence clustering, the first consideration is what kind of data should be used for clustering. We found that the direct use of raw signal information combined with the DTW algorithm can produce good clustering performance, but the time cost is high. Using the read information combined with existing clustering tools is fast but cannot produce good clustering performance. The hybrid clustering algorithm proposed in this paper makes use of these two types of data for clustering. In the initial clustering stage, the read information is used to generate the initial clustering results, and in the cluster merging and refinement stages, the raw signal information is used to continuously refine the initial clustering results. From the experimental results of simulated datasets and real datasets, the clustering accuracy of the hybrid clustering algorithm is obviously better than that of various classic clustering tools. Although the speed of hybrid clustering algorithm lags behind that of the other clustering tools, the key DTW distance comparison has been accelerated by GPU and the clustering time is completely acceptable. In general, hybrid clustering algorithm is a simple, reliable and practically useful clustering algorithm.

## 4 Conclusion

This paper presents a hybrid unsupervised approach for accurate short read clustering and barcoded sample demultiplexing in nanopore sequencing. Our approach classifies the barcoded reads in a human-understandable way, starting with a consensus read generation by strict ‘base-space’ clustering and ending with a ‘signal-similarity’ based class merging and refinement. The proposed approach can report *>* 90% clustering accuracy, achieving comparable accuracy in barcoded sample de-multiplexing compared with the supervised-learning-based methods. In addition, our method is easy to use and do not need the preparation of training data. Specifically, the introducing of GPU-acceleration significantly reduce the execution time of signal similarity comparison, which makes the processing of a huge number of data possible.

## 5 Methods

### 5.1 Overview

Motivated by the above observation, we designed an unsupervised hybrid approach to achieve accurate and efficient short read clustering for Nanopore sequencing, in which the nucleobase-based greedy algorithm is utilized to obtain initial clusters, and the raw signal information is measured to guide the continuously optimization and refinement of clustering results. Figure 7 shows the detailed workflow of hybrid clustering.

**Figure 7.**
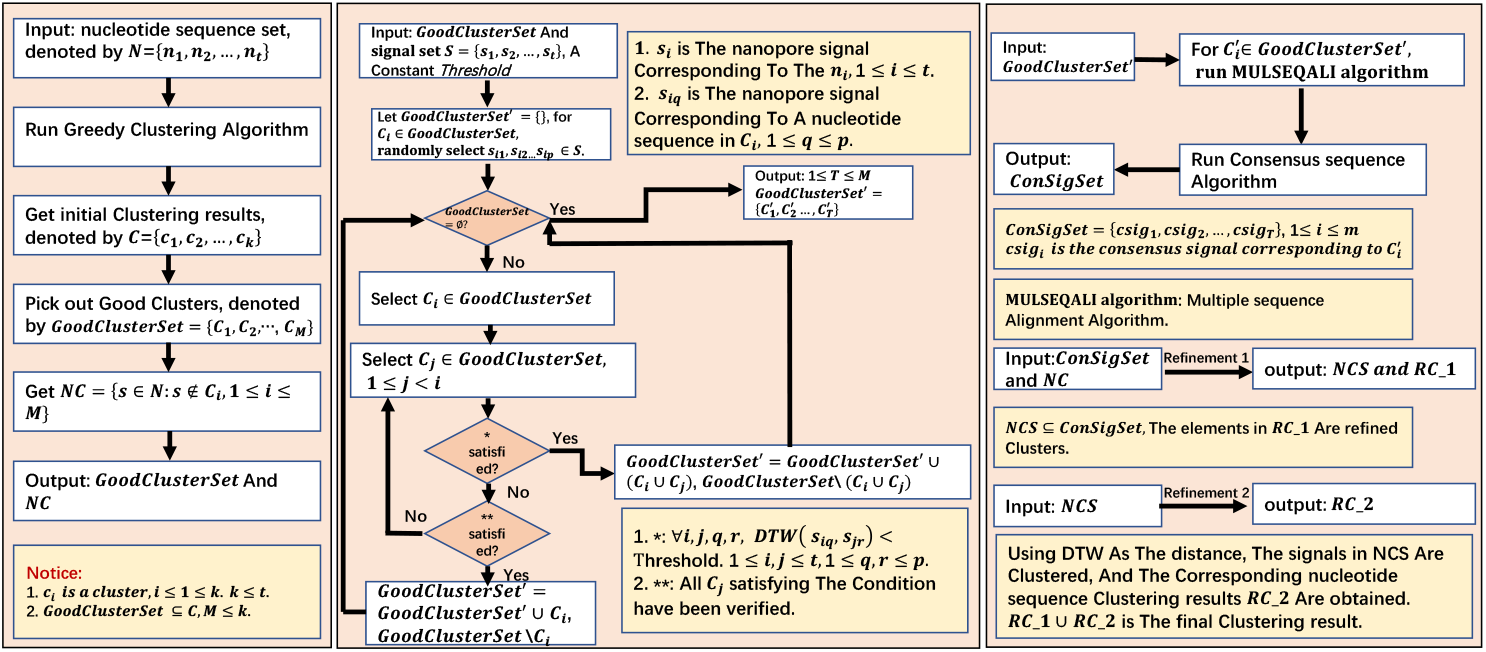
The workflow of hybrid clustering. First, a greedy clustering algorithm is used to obtain initial clustering based on the nucleotide information. Then, we try to merge the initial clusters with good homogeneity by the information of raw signals, in which the threshold of signal-to-signal DTW distance is determined with a partial sampling method. For each merged cluster, a corresponding consensus signal is produced by considering all the raw signal information within the cluster. Finally, based on these consensus signals, a signal-similarity based classification strategy is designed to assign the unclassified sequences to the already known clusters, to further refine the clustering result.

Given the nanopore sequences, we first utilize the nucleotide information for **Initial clustering** to generate clusters with high homogeneity (identity ⩾85%), whose process is based on a greedy clustering strategy and very quick. Then, we select partial sequences and rerun the initial clustering on the selected partial dataset with identity ⩾90% to observe the homogeneity of raw signals for each cluster, with which the **Threshold determination** is carried out for signal-similarity based cluster merging. Finally, we make the **Cluster merging and refinement** by calculating the DTW distance between the raw signals and each cluster’s representative signals, with **GPU-accelerated DTW** to ensure runtime. In the following, we give out the details of each step in the hybrid clustering, where the detailed pseudocode for each step is given in Supplementary Material S3.

### 5.2 Initial clustering

We utilize a nucleobase-based greedy algorithm to generate homogeneous initial clustering, in which the process is similar to the ones in CD-HIT. Figure 8 describes the detailed workflow. Firstly, the sequences are sorted in descending order of the sequence length. The longest sequence is assumed to be the representative sequence of a cluster. A short word filter[13] is applied to reduce the comparison in pairwise alignment. Here, each selected sequence is compared with the existing representative sequences. If the similarity between the selected sequence and a representative sequence is higher than the threshold, the selected sequence will be merged into the cluster of the representative sequence. Otherwise, the selected sequence becomes a new representative sequence. Repeating this process until all the sequences are visited, resulting in a number of clusters and a set of ultra-short sequences that do not belong to any cluster. Finally, A verification mechanism is used to check the homogeneity of the clusters and retrieve the ultra-short sequences which are misclassified.

**Figure 8.**
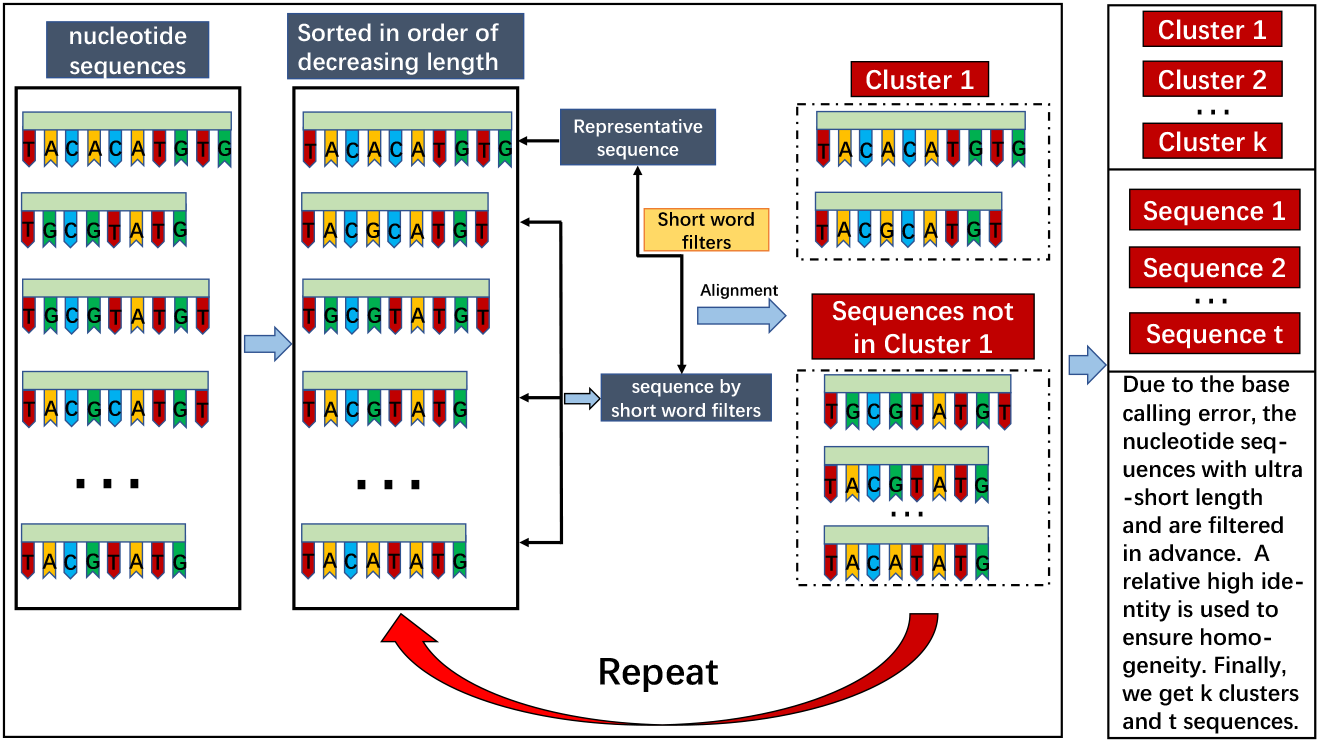
The workflow of initial clustering.

With a high enough threshold (e.g., identity ⩾85%), the initial clustering is able to quickly generate clusters with high homogeneity, where these initial clusters can be considered as completely correct, to significantly reduce the pair comparison in further signal-similarity based clustering.

### 5.3 Threshold determination

After obtaining the initial clustering result, we need to further refine it according to the raw signal information. The refinement of the initial clusters depends on the merging threshold, which is critical to the final demultiplexing accuracy. Because of the initial clusters’ high homogeneity, the merge threshold is possible to be determined from the initial clustering result. The specific process of determining the merge threshold is as follows: First, randomly select a part of the input dataset and run the initial clustering algorithm with high identity (90%) to obtain a partially initial clustering. Then, choose the largest cluster in the clustering result (denoted as *Cluster*_*max*_), get the corresponding raw signal sequences in *Cluster*_*max*_, and calculate the pairwise DTW distance of these raw signal sequences. Finally, calculate the average value of all DTW distances (represented by *dist*_*ave*_), and define *threshold* = *dist*_*ave*_ + *seq_len* + *c*, where *c* is a constant, *seq_len* indicates the length of the sequence. Here, the value of *c* can be empirically determined by a large number of simulation, the value of *seq_len* is determined by the average lengths of -nucleotide sequences in the dataset.

### 5.4 Cluster merging and refinement

#### Cluster merging

For a certain multiplex sequencing configuration, it is possible to estimate the minimal set size of a cluster. Here, we define an initial cluster with set size larger than *GoodIndex* as a good cluster, and denote *GoodClusterSet* = {*C*_1_, *C*_2_, …, *C*_*M*_ } as the set of good clusters of the initial clustering result, where *M* is the number of good clusters and {*C*_*i*_} is sorted in the descending order according to their cardinality.

For each *C*_*i*_, we randomly select *K* raw signal sequences within *C*_*i*_, and record these signals as 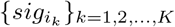, where *K* must satisfy

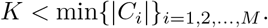

Every time we choose the top unvisited cluster *C*_*i*_, i.e., the largest unvisited cluster in *GoodClusterSet*, as a query to compare with the other clusters (*i* = 1 in the first time). We compare 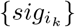 with the sampled *K* raw signals 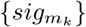 from the rest clusters (1 *< m < M*) by the DTW distance. If for 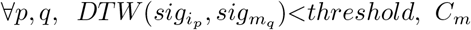 is merged into *C*_*i*_, and *GoodClusterSet* = *GoodClusterSet*\*C*_*m*_. Every time, the selected cluster *C*_*i*_ is compared with the remaining clusters {*C*_*m*_} and is merged with all the clusters *C*_*m*_ that satisfy the DTW distance constraint. We iteratively select the top unvisited cluster and make the cluster merging until all the clusters’ relationship has been checked.

#### Refinement1

It should be noted that the *GoodClusterSet* has been changed during cluster merging. After the final merging, a set of refined clusters can be obtained, i.e., 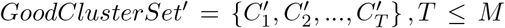. For each 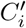, a multiple sequence alignment algorithm is possible to be applied on the (nucleotide) sequences and generate a representative consensus sequence (Figure S3 in Supplementary Material S3). The corresponding consensus sequence set can be denoted as

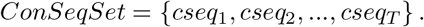

Given the consensus sequences, a standard signal can be inferred from the 6-mer table provided by Nanopore Technologies [37]. Thus, with the consensus sequences, a set of consensus signal can be generated, which is denoted as

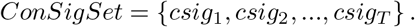

The consensus signal is utilized as standard reference to optimize the initial clustering results. For the sequences that are not in *C*_*i*_, *i* = 1, 2, …, *M*, we get the corresponding raw signals of these sequences and calculate the DTW distance between these sequences and the consensus signals in *ConSigSet*. For a given sequence, if the distance between this sequence’s raw signal and a consensus signal is less than the *threshold*, the sequence is merged to the consensus signal’s corresponding cluster.

#### Refinement2

After the above steps, there are still some sequences that have not been classified. We get the raw signals of these sequences and make the following process: First, randomly select a sequence to generate a new cluster *C*_*new*_, where the selected sequence is the representative sequence of *C*_*new*_, denoted by *seq*_*new*_. Calculate the DTW distance between the raw signal of *seq*_*new*_ and the raw signal of the remaining sequences. If the distance is less than *threshold*, add the corresponding sequence to *C*_*new*_. Repeat the process until all the sequences are visited.

Figure 1 illustrates an example for the merging and refinement process of 30 sequences from two cells. The clustering accuracy is continuously improved with the utilization of all the information residing in the nucleobase sequences and raw signals.

### 5.5 GPU-accelerated DTW

The most computational expensive part of our approach is the calculation of the tens of millions to hundreds of millions of DTW distances. Generally, the computational complexity of a DTW algorithm should be *O*(*mn*) if the algorithm is sequentially implemented, where *m* and *n* are the lengths of the compared sequences. However, with the development of graphics processing unit (GPU) for general purpose processing, CUDA (or Compute Unified Device Architecture) has been widely used to accelerate computational biology tasks [38, 39, 40]. Here, we propose a CUDA-based GPU-accelerated DTW to solve the speed problem, by combining a coarse-grained block-wise acceleration strategy and a fine-grained multi-thread acceleration strategy. Figure 9 describes the outline of our acceleration strategy.

**Figure 9.**
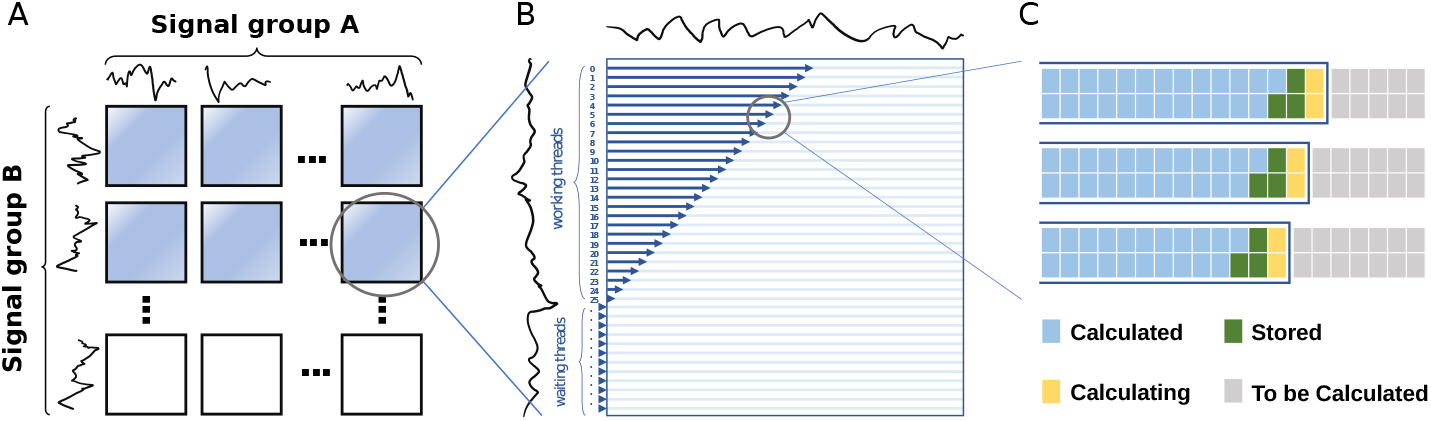
GPU-supported parallelization strategy. (A) Different signal pairs’ DTW distances are computed simultaneously within a CUDA block. (B) Each DTW matrix is calculated with multi-threads, where a register variable is used to control the wait relationships of all threads. (C) A thread can simultaneously compute multiple rows, where the values needed for the yellow cell (to be computed) are already stored in the shared memory (green cell).

#### Dependency analysis

Obviously, the DTW distance between different signals is totally independent. Thus, a coarse-grained block-wise parallel strategy is devised to calculate these DTW distances simultaneously within each CUDA block, as shown in Figure 9A. The computation of DTW weight matrix could also be accelerated by CUDA. However, data dependence exists in the calculation of a DTW matrix, i.e., the calculation of position (*i, j*) in the DTW matrix needs the values in position (*i* − 1, *j*), (*i* − 1, *j* − 1) and (*i, j* − 1). Here, considering the elements on a slash lane of a DTW matrix, these elements are independent with each other (Figure 10). Thus, we change the calculation of DTW matrix from sequence order into slash-lane order, and propose a fine-grained multi-thread parallel strategy to ensure the speed and accuracy, as shown in Figure 9B.

**Figure 10.**
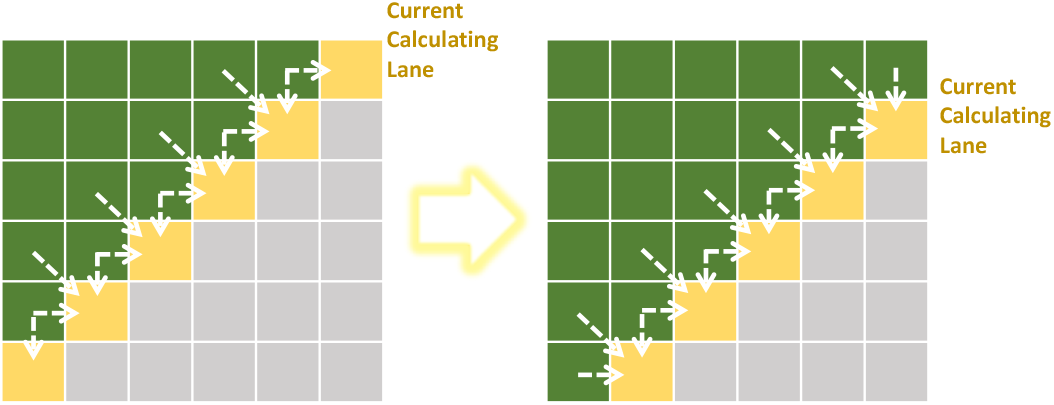
Data dependence within the calculation of a DTW matrix. The white dash arrow shows the dependence of element values if a certain element is to be calculated. Nevertheless, all the yellow elements have no dependence with each other, which enables their parallel calculation.

#### Block-wise acceleration

Within a general GPU card with NVIDIA Turing architecture, up to a few million blocks are allowed to execute asynchronously and concurrently. As shown in Figure 9A, each CUDA block is responsible for calculating one DTW distance. That is, millions of blocks could be initialized to calculate these DTW distances simultaneously, which makes the calculation extremely fast. In contrast, a multi-CPU server may only contain a few dozen cores, allowing the simultaneous calculation of only a few dozen DTW distances.

#### Multi-thread acceleration

Each DTW matrix is calculated by multiple threads lane by lane. Synchronize strategy is applied to ensure that the values needed by the current position have been calculated correctly. To control which columns should be calculated at a given time, we use a register variable *T* to serve as a timer. ∀*i* ∈ {0, 1, 2, …, *n* − 1}, the *i*th thread calculates the *i*th row (counting from 0), then the thread with thread number *t* needs to process the (*T* − *t*)th element of the row at time *T*. And the threads with thread number *c* = *T* − *t <* 0 should wait in place until *T* − *c >* 0. Figure 9B shows an example with *T* = 25.

#### On-chip storage

Since a CUDA block contains 1024 threads at most but the longest signal length is up to ∼1500 (a barcode’s length is up to 145, while the corresponding signal is 8 ∼ 10 times of the barcode sequence), we extended the algorithm to let a thread computes two DTW matrix rows at a time, which makes a block able to process 2048-length signal. Considering barcode sequences are not too long (such as 145 bp), the GPU card of the current Turing architecture can fully store data and perform calculations in on-chip memory (shared memory), which avoids the copy cost from the global memory and further accelerates the calculation. As shown in Figure 9C, green cells represent the elements stored in shared memory, and the yellow cells are the elements being calculating. Actually, maximum amount of shared memory per block is 163KB on NVIDIA Turing architecture, which provides the ability that one thread processes 9 DTW rows whose elements stored in single-precision float format.

## 6 Acknowledgements

We are grateful to Prof. Lei M. Li for the discussion and suggestion of methods and experiments.

## 8 Availability of data and materials

The data and code is available at https://github.com/junhaiqi/Hybrid_clustering.git.

## 10 Authors’ contributions

G.L., X. G. and F. Z. conceived and managed the project. R.H. and J. Q. implemented the algorithm, collected all the datasets, and performed all the analysis. Y.X. and X.S. were involved in data analysis and testing of the algorithm. All authors have read and approved the final manuscript.

